# Chimeric music reveals an interaction of pitch and time in electrophysiological signatures of music encoding

**DOI:** 10.1101/2024.11.04.621689

**Authors:** Tong Shan, Edmund C. Lalor, Ross K. Maddox

**Affiliations:** Department of Biomedical Engineering, University of Rochester, Rochester, NY, USA 14627; Department of Neuroscience, University of Rochester, Rochester, NY, USA 14627; Del Monte Institute for Neuroscience, University of Rochester, Rochester, NY, USA 14627; Kresge Hearing Research Institute, Department of Otolaryngology-Head & Neck Surgery, University of Michigan, Ann Arbor, MI, USA 48109

## Abstract

Pitch and time are the essential dimensions defining musical melody. Recent electrophysiological studies have explored the neural encoding of musical pitch and time by leveraging probabilistic models of their sequences, but few have studied how the features might interact. This study examines these interactions by introducing “chimeric music,” which pairs two distinct melodies, and exchanges their pitch contours and note onset-times to create two new melodies, thereby distorting musical pattern while maintaining the marginal statistics of the original pieces’ pitch and temporal sequences. Through this manipulation, we aimed to dissect the music processing and the interaction between pitch and time. Employing the temporal response function (TRF) framework, we analyzed the neural encoding of melodic expectation and musical downbeats in participants with varying levels of musical training. Our findings revealed differences in the encoding of melodic expectation between original and chimeric stimuli in both dimensions, with a significant impact of musical experience. This suggests that the structural violation due to decoupling the pitch and temporal structure affect expectation processing. In our analysis of downbeat encoding, we found an enhanced neural response when participants heard a note that aligned with the downbeat during music listening. In chimeric music, responses to downbeats were larger when the note was also a downbeat in the original music that provided the pitch sequence, indicating an effect of pitch structure on beat perception. This study advances our understanding of the neural underpinnings of music, emphasizing the significance of pitch-time interaction in the neural encoding of music.

**Significance Statement:** Listening to music is a complex and multidimensional auditory experience. Recent studies have investigated the neural encoding of pitch and timing sequences in musical structure, but they have been studied independently. This study addresses the gap in understanding of how the interaction between pitch and time affects their encoding. By introducing “chimeric music,” which decouples these dimensions in melodies, we investigate how this interaction influences the neural activities using EEG. Leveraging and the temporal response function (TRF) framework, we found that structural violations in pitch-time interactions impact musical expectation processing and beat perception. These results advance our knowledge of how the brain processes complex auditory stimuli like music, underscoring the critical role of pitch and time interactions in music perception.

## Introduction

Pitch and time are the two fundamental dimensions of music that together form the complex structure and organized sequence of events, forming the basis of a melody (Lerdahl, 2019; Lerdahl & Jackendoff, 1996; Tillmann, 2012). The interaction between these dimensions can influence the perception of each other (Krumhansl, 2000; Prince, 2011; Prince et al., 2009). For example, the detection of pitch deviants depends on temporal structure regularities (Herbst & Obleser, 2019; Jones et al., 1982; Jones et al., 2002). Rhythm processing is also affected by the tonal structure and the predictability of pitch (Keitel et al., 2023; Tillmann & Bharucha, 2002). This interplay exemplifies the complexity of music as a multidimensional auditory experience.

When processing music, listeners’ experience with contextually relevant musical structure (i.e., the underlying syntax) will be activated, from which expectancy emerges, influencing the processing of subsequent events (Huron, 2008; Tillmann, 2012). Neuroimaging studies have shown that violating expectations triggers neural responses to unexpected musical patterns (Koelsch, 2009; Koelsch et al., 2000; Vuust et al., 2012).

Recently, temporal response functions (TRF) have been used to study the neural tracking of continuous stimuli by modeling ongoing brain activity based on specific stimulus features (Crosse et al., 2016; Ding & Simon, 2012). Using this method, studies have provided insights on the neural tracking of melodic expectations in continuous music (Bianco et al., 2024; Di Liberto, Pelofi, Bianco, et al., 2020; Kern et al., 2022; Marion et al., 2021). These investigations reveal that the melodic expectation process is inherently active while humans listen to music, with quantified features like musical note surprise and uncertainty being key components. These components are computed from probabilistic models, with one such model, the Information Dynamics of Music (IDyOM) (Pearce & Wiggins, 2012; Pearce, 2005) seeing frequent use. This model computes expectation for both pitch and time, but does not consider their interaction, leaving a gap in understanding of how that interaction contributes to the neural encoding of musical expectations.

Another aspect of music processing involves meter perception, reflecting perceived temporal regularity within music (Grahn, 2012; Snyder & Krumhansl, 2001) at multiple scales. This temporal structure perception also establishes an expectancy that the events occurring on strong beats (e.g., downbeats that starts a measure) are processed as more salient than those “off the beat” (Bouwer et al., 2014; Fitzroy & Sanders, 2015; Geiser et al., 2010; Moon et al., 2020; Schaefer et al., 2011; Tierney & Kraus, 2013, 2015). However, few studies have explored how pitch’s structure influences this temporal structure perception, and those that exist relied on repetitive, simple music excerpts rather than continuous melodies (Lelo-de-Larrea-Mancera et al., 2017; Zhang et al., 2019).

To better understand the interaction of pitch and time in music listening, this study introduces a new type of auditory stimulus: chimeric music (**Figure 1a**), which exchanges the pitch and rhythm structures from different natural music pieces, resulting in new compositions that sound less structured and predictable. By decoupling the pitch and time dimensions and violating their traditional linkage while preserving the marginal statistics of the original pieces, it provides a way to study pitch-time interaction.

**Figure 1.**
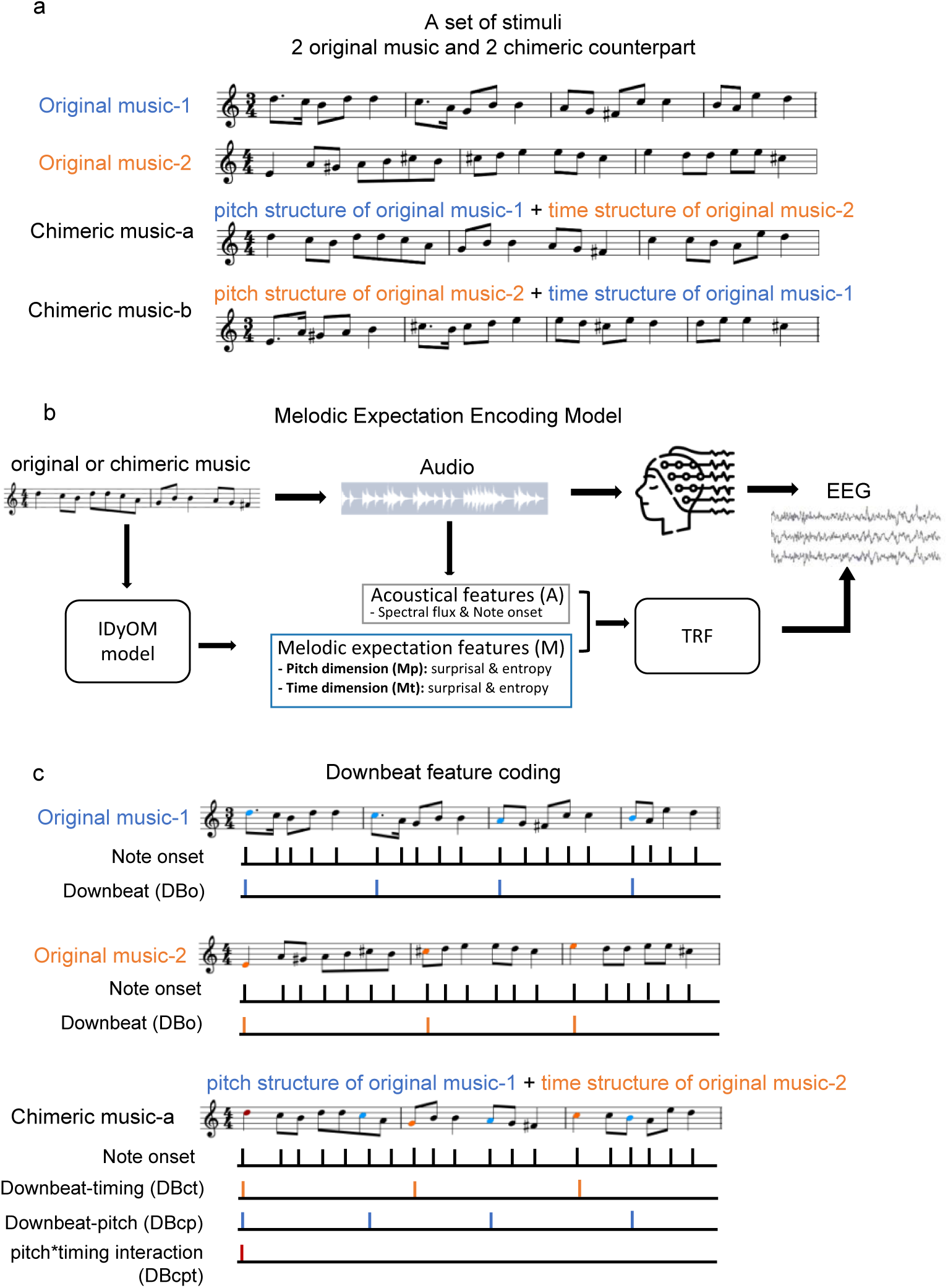
Overview of the research framework. (**a**) An example set of stimuli consists of 2 original music and their 2 chimeric music counterparts. The chimeric music-a has the pitch structure of original music-1 and time (rhythm) structure of original music-2. The chimeric music-b has the pitch structure of original music-2 and time structure of original music-1. (**b**) TRF paradigm for the melodic expectation encoding. Participants listen to the monophonic music, including original and chimeric music. Acoustical features including spectral flux and note onsets were computed from the music. The IDyOM model was used to calculate the note-level melodic expectation features for pitch and time, including pitch surprisal, pitch entropy, time surprisal, and time entropy. The acoustical features (A) and the melodic expectation features (M) were used as the regressors in the TRF model. (**c**) The regressor illustration for downbeat encoding model. For original music, binary-coded impulse trains served as a regressor that indicate when the note onset and downbeat (DBo) occur. For chimeric music-a, in addition to the note onset, the downbeat-time (DBct) was inherited from the original music-2 downbeat regressor (marked orange), whereas the downbeat-pitch (DBcp) was from the original music-1 downbeat regressor (marked blue). The interaction term (DBcpt) was when the note is both DBct and DBcp (marked red).

Our initial investigation focused on the neural encoding of melodic expectation within the TRF framework (**Figure 1b**), following the methodologies of previous studies (Bianco et al., 2024; Di Liberto, Pelofi, Bianco, et al., 2020; Kern et al., 2022). We compared the neural tracking of the original music and their chimeric counterparts. Secondly, we studied the influence of pitch structure on the time structure through downbeat encoding. The downbeat, defined as the first beat of a measure and typically the strongest metrical accent (Jehan, 2005), serves as a focal point for this analysis. We used a TRF framework alongside a set of binary-coded indicator features according to the note onset and metrical structure (**Figure 1c**). We found both the neural encoding of melodic expectation and musical downbeats were influenced by pitch-time interactions in musical structure.

## Methods and Materials

### Participants

Twenty-seven subjects participated in this experiment with an age of 22.9 ± 3.9 (mean ± STD) years. All participants had audiometric thresholds of 20 dB HL or better from 500 Hz to 8000 Hz. Participants self-reported having normal or corrected to normal vision. Following the experimental session, subjects completed a self-reported musicianship survey (see appendix for the survey).

All subjects gave informed consent and received compensation for their time. Data collection was conducted under a protocol approved by the University of Rochester Subjects Review Board (#66988).

### Original Music and Chimeric Music Stimuli

Monophonic European folksongs from the Essen Folksong Collection (Schaffrath, 1995) corpus were used as the stimuli in this study. We selected pieces that had durations longer than 25 seconds, excluding those with less discernable melody lines or many repetitions. The selected pieces were then converted to MIDI, a format which encodes the note value, onset, and duration of each note.

To explore the interaction between musical pitch and time, we created a new stimulus called “chimeric music”. Note pitch sequence and the note onset times sequence were extracted from each piece using python package *Mido* (Bjørndalen & Doursenaud, 2023). Chimeric music was created by combining the pitch sequence from one song with the note time sequence of another, forming pairs as illustrated in **Figure 1a**. Pairs were made on the following approach: music pieces were first categorized into two beat structure groups based on their time signatures – either triple beats (3/4, 3/8, and 6/8 time signatures) or duple / quadruple beats (2/4, 4/4, and 4/8 time signatures). Pieces were then paired such that each pair consisted of one from each beat group. When fusing the pitch and time, the note matching was always starting with the beginning ensuring the same context prior in each domain. To avoid significant duration loss, the paired pieces had close to the same number of notes, so that the synthesized chimeric music had similar duration to the original. This pairing strategy was designed to disrupt the structured melodic patterns by combining pitch contours associated with triple time signatures with duple or quadruple beat timing, or vice versa.

Each stimulus set contained 2 original matched music and 2 chimeric music counterparts. In total, 33 sets of stimuli were present in this study, including 66 original music and their 66 chimeric music counterparts. Both original and chimeric music stimuli were rendered into WAV files by *FluidSynth* using its default acoustic grand piano SoundFont with a sampling rate of 48 kHz. All music notes in all music maintained uniform velocity (i.e., same intensity), and the overall amplitudes of the stimuli were normalized to a root mean square of 0.01, ensuring consistent loudness across stimuli.

### Stimulus Presentation and Procedure

During the experimental session, subjects were seated in a sound-isolating booth facing a 24- inch BenQ monitor, where they passively listened to the provided stimuli. Each stimulus was presented at a consistent average level of 60 dB SPL. The stimulus audio was delivered through ER-2 insert earphones (Etymotic Research, Elk Grove, IL) connected to an RME Babyface Pro digital sound card (RME, Haimhausen, Germany). The stimulus presentation for the experiment was controlled by a python script using the *expyfun* package (Larson et al., 2014). There were in total 132 music trials with duration ranging from 27 seconds to 50 seconds, resulting in a total duration of the experiment around 77 minutes. The stimuli were played in random order to subjects. Following each music trial, subjects were asked to rate their preference on the monitor for the heard piece on a scale of 1 through 7, with 1 indicating strongly disliked and 7 strongly liked.

### EEG Data Acquisition and Preprocessing

Subcortical and cortical EEG signals were simultaneously recorded using the ReCorder software from BrainVision (Brain Products GmbH, Gilching, Germany). For the acquisition of subcortical signals, passive Ag/AgCl electrodes were placed frontocentrally (FCz, non-inverting), left and right earlobes (A1, A2, inverting references), and the frontal pole (Fpz, ground). These electrodes were connected to two EP-Preamp amplifiers connected to an actiCHamp Plus recording system (both from Brain Products GmbH, Gilching, Germany). For the acquisition of cortical signals, a 32-channel configuration was employed using active electrodes, arranged according to the International 10–20 system. The 32-channel system was directly connected to the ActiCHamp plus. To ensure good signal quality, the impedance of all electrodes was verified to be below 10 kΩ before starting the recordings. Both signals were recorded at a high sampling rate of 10k Hz.

The subcortical signal was high-pass filtered offline at 1 Hz using a first-order causal Butterworth filter to remove the slow drift. We also used a second-order infinite impulse response notch filter to remove 60 Hz line noise and its multiples with a width of 5 Hz (Shan et al., 2024). The left and right channels were averaged as the final subcortical EEG signal. The cortical signal was high-passed at 1 Hz and low-passed at 8 Hz with bidirectional zero-phase FIR filter and downsampled to 125 Hz. All channels were re-referenced to the average of the two mastoid channels TP9 and TP10 to highlight the auditory responses (Luck, 2014). To repair the eye movement and blink artifact, an Independent Components Analysis (ICA) was conducted using *MNE* package (Ablin et al., 2018; Gramfort et al., 2013; Larson et al., 2023), and epochs with excessive variance were excluded from subsequent analysis.

### Cortical EEG Analysis

The temporal response function (TRF) (Crosse et al., 2016; Ding et al., 2014; Lalor et al., 2009), a time-resolved regression analysis technique, was employed in this study to compute the neural tracking of music features using the mTRF toolbox (Crosse et al., 2016) and its adapted python version (Bialas et al., 2023). The TRF has been widely used to map the cortical neural activity evoked by continuous auditory stimuli, such as speech and music (Crosse et al., 2016; Di Liberto et al., 2015; Di Liberto, Pelofi, Bianco, et al., 2020; Ding et al., 2014). The TRF was computed by a linear regression to estimate model weights that optimally represent the relationship between neural activity and single or multiple stimulus features. The derived TRF function acts as a filter that is interpretable in sensor space and time lag, offering insights comparable to those provided by event-related potentials (ERPs).

For this study, we aimed to explore different aspects of neural tracking of music by fitting two distinct groups of TRF models. These models were designed to incorporate various groups of features, as detailed in the subsequent sections.

### Melodic Expectation Encoding

The first aspect we explored was the melodic expectation encoding in original and chimeric music and their differences. The expectation features were note-level markers derived from the Information Dynamics of Music (IDyOM) model (Pearce & Wiggins, 2012; Pearce, 2005), which is a hidden Markov model that learns the stimulus statistical patterns and predicts the probabilistic structure of symbolic sequences within specific stimulus domains. For a given musical context, IDyOM estimates the likelihood of the next musical note based on the prior *n* − 1 sequence (i.e., n-gram context). The model gives the output of the sequences in two aspects: 1) Surprisal (*S*) or information content, reflecting the unexpectedness of the current note *x*_*t*_ at time *t* given the preceding notes 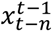, calculated as

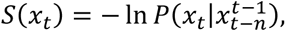

and 2) the Entropy (*E*), reflecting the uncertainty of the previous context before the *x*_*t*_ happens, computed across all notes in the set of possible notes (*x* ∈ *K*) as follows

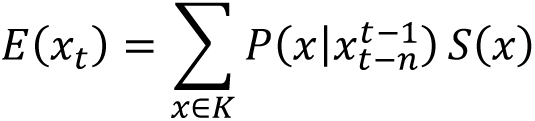

These computations were derived from different musical dimensions, or viewpoints. In this study, we focused on the pitch (IDyOM viewpoint *cpitch*) and inter-onset interval (IDyOM viewpoint *ioi*) to analyze the pitch and time dimensions, denoted as Sp (pitch surprisal), Ep (pitch entropy), St (time surprisal) and Et (time entropy).

The IDyOM model can incorporate both a short-term model (STM) and long-term model (LTM), where the STM is trained on the current stimulus trial and the LTM is trained on broader corpus material simulating statistical learning through life-time musical enculturation. In this study, we used melodic expectation features derived from a combination of STM and LTM (termed “both” model in IDyOM), in line with previous work (Di Liberto, Pelofi, Bianco, et al., 2020). The LTM was trained on European folksongs also from Essen Folksong Collection (Schaffrath, 1995) corpus excluding the songs that were used as stimuli.

Following the previous studies investigating the melodic expectation encoding in music (Bianco et al., 2024; Di Liberto, Pelofi, Bianco, et al., 2020; Kern et al., 2022), an acoustical model (denoted as A) was fitted with continuous spectral flux and the note onset (a binary indicator with impulses where a note onset occurred as the acoustical regressors, which served as a baseline model. We then extended this model forming the melodic expectation TRF model (denoted as AM), which incorporated both the aforementioned acoustic features and the expectation features derived from IDyOM (**Figure 1b**). To explore the contribution of melodic expectation features from the pitch and time dimensions individually, we also tested two reduced models from the full AM model including only pitch dimension features (Sp/Ep) or time dimension features (St/Et), denoted as AMp and AMt, respectively.

Notably, given that the sequence of the pitch and time were consistent across both original and chimeric music (i.e., same context in each dimension), the resultant expectation features Sp/Ep and St/Et were identical. This design allowed TRFs to be fit using the same feature vectors for both stimulus types, with potential TRF differences indicating an effect of the pitch-time decoupling inherent to the chimeric stimuli.

Ridge regression was used to train the TRF models to avoid overfitting (lambda values ranges from 10^-3^ to 10^3^). We used leave-one-out cross-validation (across trials) to evaluate the TRF models on unseen data. The lambda for each TRF model and for each subject providing the best prediction of unseen data was chosen. The average Pearson’s correlation between the predicted EEG signal and the recorded real EEG signal across all channels was used as the metric to assess the TRF model’s prediction accuracy.

### Music Downbeat Encoding

The second aspect we explored was downbeat encoding in original and chimeric music. Initially, we established a note onset regressor, made as an impulse train where each musical note occurrence was marked by a unit impulse. For the original music pieces, a downbeat regressor (DBo) was similarly created as an impulse train, coding the onsets of downbeat notes as 1 and all other timepoints as 0 (the non-zero points in this regressor were thus a subset of the onset regressor). In the case of chimeric music, we delineated two types of downbeats: downbeat-pitch (DBcp) represented notes that were originally downbeats in the piece whose pitch contour was included in the chimeric stimulus, and downbeat-time (DBct) represented notes that were downbeats in the piece whose rhythmic sequence was used. Note that the downbeat-time notes were also the downbeat note by definition for the chimeric stimulus, as downbeats are temporally defined. We also created a pitch*time interaction regressor (DBcpt) as an impulse train indicating the note is simultaneously a downbeat-pitch note and downbeat-time note. An example of these regressors is shown in **Figure 1c**. Chimeric music-A has the pitch contour of original music-1 and rhythm contour of original music-2. The 6^th^ note (C5 highlighted with blue) was marked as downbeat-pitch originating from original music-1, while the 8^th^ note (G4 highlighted with orange) marked as downbeat-time originating from original music-2.

In the downbeat encoding analysis, we were interested in comparing neural encoding through the TRF weights for each of the downbeat regressors. Therefore, we trained TRF models from all the trials for original or chimeric music for each subject. Since the regressors were all impulse trains, we used the ordinary least squares (OLS) regression instead of ridge regression in order to preserve the interpretability of the TRF weights.

## Statistical Analysis

### Melodic expectation encoding TRF analysis

For the melodic expectation encoding TRF analysis, a linear mixed-effect model was applied to analyze the prediction accuracy (measured by the correlation, Pearson’s R) with the TRF model types (A or AM), the stimulus category (original or chimeric music), the musicianship (subjects’ formal music training years), the preference ratings and their interaction. These factors were considered as fixed effects, while individual subjects and the stimulus set were treated as random effects. The model formula used was

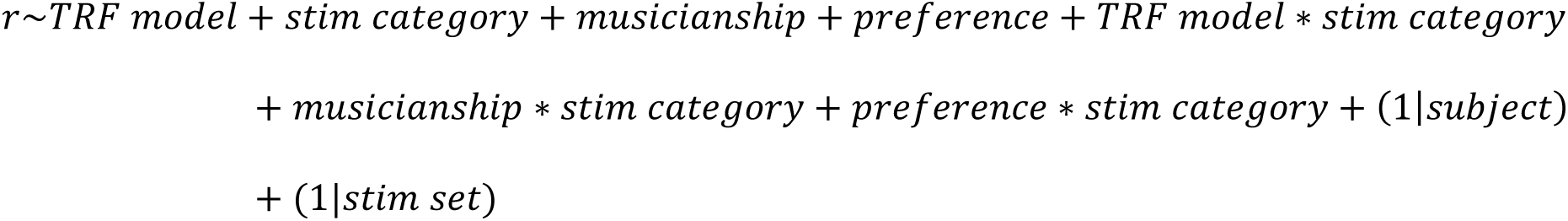

To further assess the enhancement effect of melodic expectation, we calculated the difference in prediction accuracy between each melodic expectation model and the acoustic baseline model within each subject and stimulus set. A subsequent linear mixed-effect model was fitted with TRF model types of contrast (AMp−A, AMt−A, AM−A), the stimulus category (original or chimeric music), the musicianship and their interaction terms as fixed effect, again considering subject and the stimulus set as random effects. The formula for this analysis was

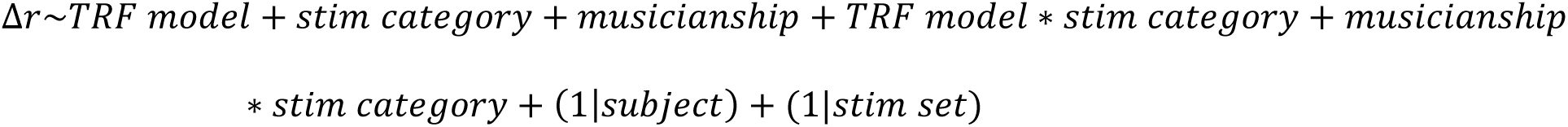

Post-hoc paired t-tests with Holm-Bonferroni correction were performed to determine the statistical differences between each model type and stimulus category.

Additionally, we visualized the derived TRF weights for each regressor, utilizing a permutation test with clustering to identify significant differences in TRF weights between original and chimeric music.

### Downbeat encoding TRF analysis

To determine downbeat encoding, we displayed the TRF weights for each regressor, employing permutation tests with clustering for statistical inference. For original music, TRF weights for note onset and DBo regressors were contrasted against zero (the null response). Similarly, for chimeric music, TRF weights for note onset, DBcp, DBct, and DBcpt were also contrasted against zero. A comparison between the note onset TRF weights from original and chimeric music was conducted, as well as the DBo and DBct TRF weights.

Pearson’s correlation analysis was also used to explore the relationship between the amplitude of derived TRF weights and the length of formal musical training (in years) of the participants, while applying false discovery rate (FDR) correction for multiple comparisons.

### Subcortical EEG Analysis

In the derivation of the subcortical auditory brainstem response (ABR), a deconvolution paradigm was used as described in Shan et al. (2024). These responses are also calculated as TRFs, but with some key differences. They were recorded with passive electrodes and special preamps suitable for the ABR (BrainProducts EP-Preamp), and analyzed at a much higher sampling rate of 10,000 samples / s. The input features were created by passing the stimuli through a model of the auditory periphery which outputs the auditory nerve firing rate. The resulting TRFs show responses at much shorter latencies that mirror the standard ABR morphology.

The grand averaged ABR waveforms from original and chimeric music were then compared. Statistical inference was determined by timepoint-wise Wilcoxon signed-rank test with FDR correction.

## Results

### Expectation effect is larger in original than in chimeric music

Our first analysis confirmed a significant preference for original music over chimeric music (p<0.001, paired-t test; **Figure 2a**).

**Figure 2.**
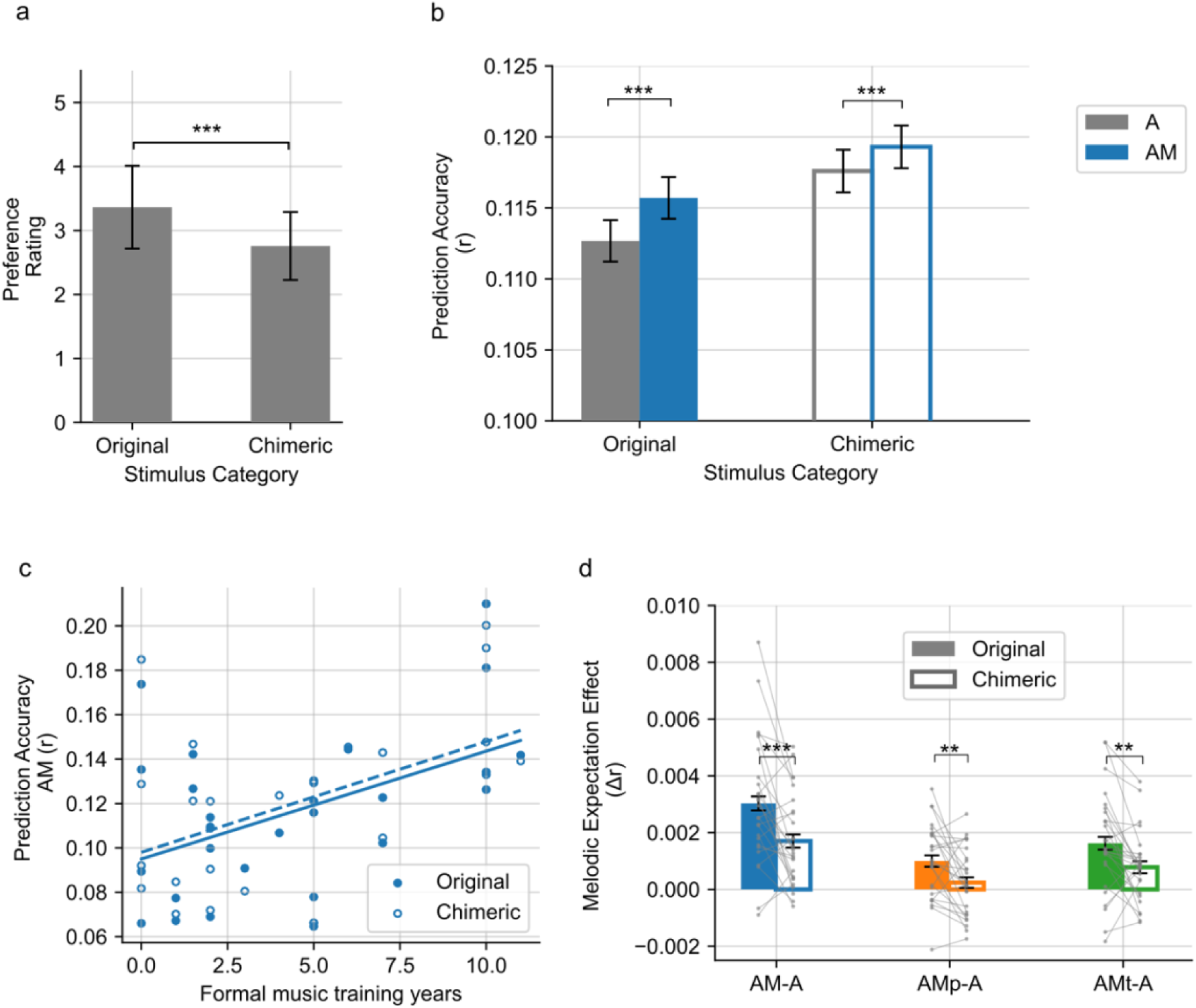
**(a)** Preference rating. The bars show the averaged rating of each stimulus category. Error bar indicates ± 1 SEM. **(b)** The TRF prediction accuracy of A and AM model. **(c)** The correlation between the TRF prediction accuracy of AM model and the music expertise as in formal music training years. **(d)** The melodic expectation effect as the prediction accuracy difference (Δr) between each melodic expectation models (AM, AMp, AMt) and the baseline acoustic model (A). Grey lines show the averaged Δr from each subject. *p<0.05, **p<0.01, ***p<0.001.

We used the TRF model to explore the neural tracking of melodic expectation during listening to the original and chimeric music. We compared the prediction accuracy of the four TRF models between the two stimulus categories. These models were acoustical feature only model (A) and model combing both acoustics and melodic expectation features from IDyOM’s model (AM) as described in the Methods and Materials section. The prediction accuracies for each trial were obtained using leave-one-out cross-validation. A linear mixed-effects model was subsequently fitted to examine the influence of stimulus category (original or chimeric music), model types (A or AM), subject’s musical training, and preference rating on prediction accuracy, as summarized in **Table 1**.

**Table 1.** The fitted mixed effect linear model results for prediction accuracy (Pearson’s r). Original music and model A are the baselines for the term category and model.

|  | Coeff | z | p |
| --- | --- | --- | --- |
| **Intercept (Original music A)** | 0.092 | 35.074 | <0.001 |
| **Chimeric Music** | 0.011 | 4.418 | <0.001 |
| **AM** | 0.003 | 2.772 | 0.006 |
| **Chimeric\* AM** | -0.001 | -0.858 | 0.391 |
| **Musicianship** | 0.005 | 13.325 | <0.001 |
| **Chimeric\*Musicianship** | -0.000 | -0.616 | 0.538 |
| **Preference** | -0.000 | -0.224 | 0.823 |
| **Chimeric\*Preference** | -0.002 | -3.348 | 0.001 |

In setting our baseline (intercept) as model A for original music, our analysis revealed significant enhancements in prediction accuracy for the TRF models incorporating both acoustic and melodic expectation features with AM terms tested significant (p<0.001). This improvement in predictive accuracy aligns with previous research (Di Liberto, Pelofi, Bianco, et al., 2020; Kern et al., 2022), demonstrating that including melodic expectation features better explains neural activity during music listening. Notably, the AM terms were significant for both original and chimeric music, as depicted in **Figure 2b**, suggesting that the expectation process is active even when the normal associations of pitch and temporal structure are violated.

Furthermore, our findings indicated that length of formal musical training is correlated with prediction accuracy in general (p<0.001 for the musicianship term). A subsequent post-hoc analysis confirmed that the musicianship is significantly correlated with the prediction accuracy for both original and chimeric music for both A and AM models as shown in **Figure 2c**. Interestingly, while preference alone did not significantly influence prediction accuracy, the interaction between preference and stimulus type exhibited a small but significant negative effect, indicating slightly stronger neural tracking of less preferred chimeric music.

To delve deeper into the melodic expectation effect, we examined the prediction accuracy difference (Δr) between the melodic expectation model (AM) and the baseline acoustic model (A), as well as the contribution of the pitch or time dimension features (AMp-A or AMt-A). The results, summarized in **Table 2**, show a significant negative coefficient for chimeric music, indicating a greater overall melodic expectation effect for original music compared to chimeric music. Post-hoc comparisons further revealed that prediction improvement offered by the melodic expectation features was significant for all AM, AMp and AMt models (**Figure 2d**). This suggests that both pitch dimension features and time dimension features contributed to the melodic expectation effect.

**Table 2.** The fitted mixed effect linear model results for the expectation effect (Δ*r*). Original music and AMp-A are the baselines for the term category and model.

|  | **Coeff** | **z** | **p** |
| --- | --- | --- | --- |
| **Intercept (Original music AM-A)** | 0.003 | 11.156 | <0.001 |
| **Chimeric Music** | -0.002 | -4.969 | <0.001 |
| **AMp-A** | -0.002 | -7.546 | <0.001 |
| **AMt-A** | -0.001 | -5.218 | <0.001 |
| **Chimeric \* AMp-A** | 0.001 | 1.488 | 0.137 |
| **Chimeric \* AMt-A** | 0.000 | 1.264 | 0.206 |
| **Musicianship** | -0.000 | -0.745 | 0.456 |
| **Chimeric \* Musicianship** | 0.000 | 1.617 | 0.106 |

When comparing TRF weight waveforms of each regressor from the AM model between original and chimeric music, no significant differences were identified between the two stimulus categories (all p>0.05, permutation test with cluster; refer to supplemental material **Figure S1**).

### Downbeat notes elicit distinct responses

We next explored whether the neural encoding of downbeats (note onsets that occur on the first beat of the measure), defined in time domain, is influenced by another music domain – pitch, leading to another aspect of the music processing with the pitch and time interaction. For the original music, we fitted the TRF with two binary-coded regressors – note onset and downbeat (DBo). The TRF weights, as shown in **Figure 3a**, revealed that the waveform for note onset exhibited typical auditory evoked potential features, including P1 and P2 peaks, which appeared at approximately 80 ms and 185 ms post-stimulus, respectively. The TRF weights for downbeat regressor also demonstrated a peak at around 185 ms and a dip at around 650 ms, across most of the scalp electrodes, as shown in **Figure 3b** (permutation test with cluster).

**Figure 3.**
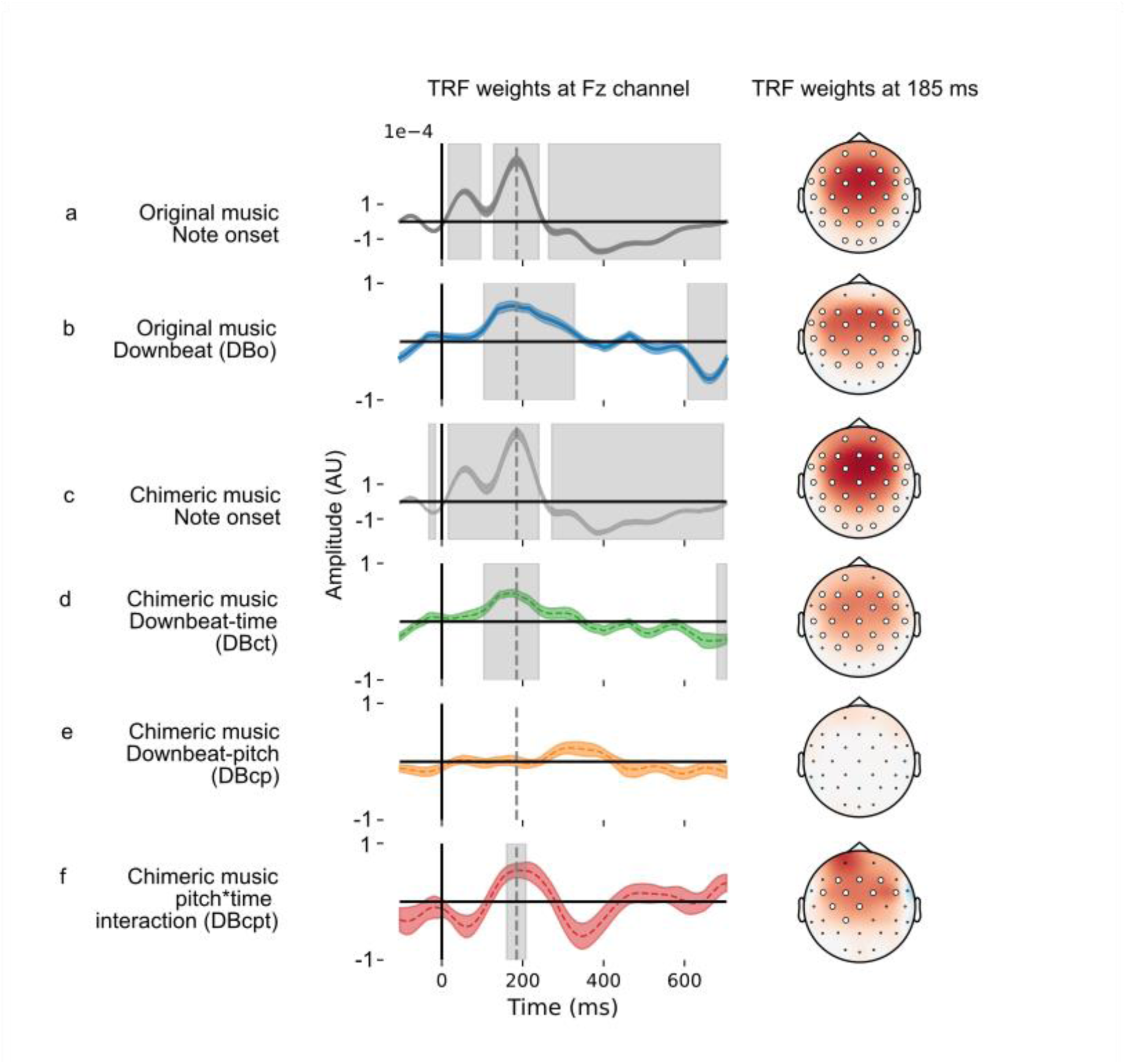
TRF weights in the downbeat encoding model for original music note onset (a), DBo (b), chimeric music note onset (c), DBct (d), DBcp (e), and DBcpt (f). Left side is the TRF weights at Fz channel. Colored shaded area shows ± 1 SEM. The grey area shows the significant time range. Right side is the topography of the TRF weighs at the peak time point, 185 ms. The larger circle shows the significant channels.

We conducted the same analysis for chimeric music, albeit with four binary-coded regressors in fitting the TRF: note onset, downbeat-time (DBct), downbeat-pitch (DBcp) and the interaction (DBcpt). Please see Methods and Materials for detail description about these regressors. The TRF weights for note onset from chimeric music shared the same morphology as that of original music (**Figure 3c**) and tested not significantly different (all p>0.05, permutation test with cluster). Similarly, the TRF weights derived from DBct mirrored those of the DBo of original music (**Figure 3d**), with no significant difference (all p>0.05, permutation test with cluster).

### Downbeat encoding is also influenced by the pitch structure

Downbeats are defined by the temporal structure of the music. But in chimeric music, we can further investigate whether pitch structure influences on this distinct downbeat neural response, via the downbeat-pitch (Dcp) and the interaction regressors (Dcpt), specifically.

The derived TRF for Dcp regressor revealed a broad peak around 260 ms on channel Fz. However, this observation did not reach statistical significance when compared to the null response, at Fz or other electrodes (**Figure 3e**; all p>0.05, permutation test with cluster). However, the TRF weights of the interaction term (Dcpt) demonstrated a peak around 180 ms (**Figure 3f**), where there was an enhancement of P2 amplitude. This enhancement is in addition to the effect of downbeat time. This result supports the hypothesis that the neural coding of beat is influenced by the pitch structure. The cluster of significant channels and timepoints of the derived TRF can be found in supplemental materials (**Figure S2-S6**).

The multivariate impulse train regressors we used in downbeat encoding models allow for an interpretation of the corresponding TRF weights like traditional ERPs waveforms. Taking chimeric music as example, when a note occurs, the TRF weight associated with a note’s onset serves as the baseline ERP. If the note is marked as downbeat-time, an additional increment represented by the TRF weight of the DBct regressor is integrated into the baseline ERP. Moreover, if the note also happens to be a downbeat-pitch, a further increment—indicated by the TRF weight of the DBcpt regressor—is added to this cumulative response. Therefore, the ERP to a note that is both a downbeat-time and downbeat-pitch is represented by the summation of TRF weights from note onset, DBct, and DBcpt. This approach incorporating binary coding regressors in TRF analysis provide a bridge between traditional ERP studies and new investigations into the neural encoding of music by disentangling complex features.

We also examined the impact of musicianship, measured by the formal music training years of the subjects, which ranged from 0 to 11 years (4.19 ± 3.58, mean ± std). To understand how musicianship influences the neural encoding of music, we used Pearson’s correlation to assess the relationship between the years of musical training and the amplitudes of TRF weights at specific peak components—P1 and P2 for note onset regressors, and P2 peak for downbeat-related regressors (DBo, DBct, and DBcpt) across all channels. Significant correlation was observed between the musical training and the maximum amplitude of the P2 component for the note onset regressor across the majority of channels (**Figure 4b** and **4d**). However, the maximum amplitude values from the TRF weights of other regressors did not exhibit a significant correlation with the musicianship (**Figure 4a, c, e, f, g, and h**).

**Figure 4.**
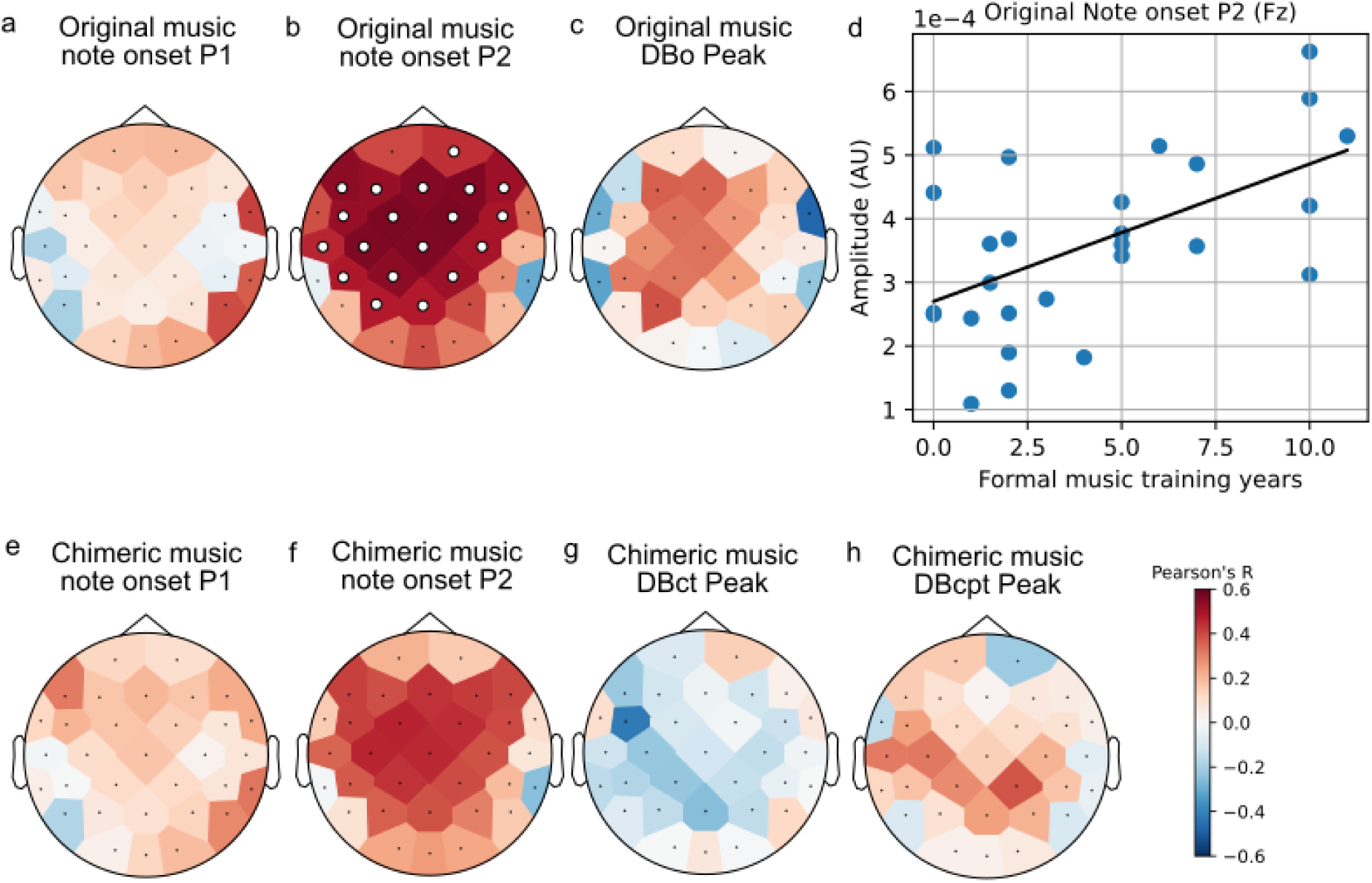
The correlation between the peak amplitude of the TRF weights for each regressor and the subjects’ formal music training years. The larger circles indicate channels that has significant correlation. For the correlation on channel Fz (d), r=0.55, p=0.015 (FDR corrected).

### Subcortical responses are same for original and chimeric music

We also investigated the ABRs as subcortical responses derived from the original and chimeric music, shown in **Figure 5**. The waveforms have the canonical ABR morphology, which is also consistent with previous study (Shan et al., 2024). The waveforms for original and chimeric music were highly correlated. No significant difference was found between the two stimulus categories (all p>0.05, timepoint-wise Wilcoxon sign rank test, FDR corrected).

**Figure 5.**
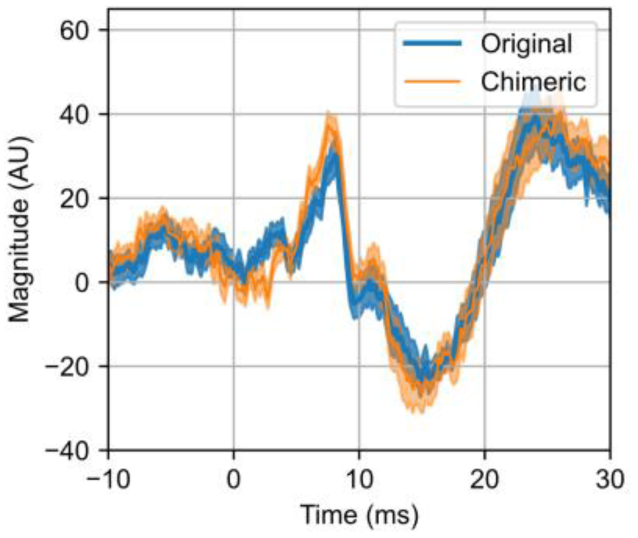
Grand averaged ABR waveforms derived from original and chimeric music. Shaded area shows ± 1 SEM.

## Discussion

By using chimeric music as a stimulus, we sought to investigate how pitch and time interact during music listening. We used TRF analyses that assess that neural tracking of continuous auditory stimuli as the main tool to study two aspects of music neural encoding: 1) melodic expectation encoding and its difference between original and chimeric music; 2) downbeat encoding and the pitch-time interaction influence on temporal structure perception.

For the melodic expectation encoding analysis, a notable advantage of using the chimeric music is that it preserved the context of each dimension, resulting in identical melodic expectation regressors to the original music while interrupting the linked role of pitch and time in building up the musical structure. Therefore, the TRF analysis and the comparison were well-controlled.

For melodic expectation we found that, consistent with previous research (Bianco et al., 2024; Di Liberto, Pelofi, Bianco, et al., 2020; Kern et al., 2022), a model that includes melodic expectation features has better predictive accuracy than the baseline acoustic-only model. This result also held true for chimeric music, even though the normal associations of pitch and temporal structure were violated (**Figure 2b**). An interesting finding was that prediction accuracy was better overall for the chimeric music. We can only guess why this was the case, but it may be related to listening more intently to atypical melodies. A previous study found that atonal melodies with the same rhythm as typical Western ones showed stronger EEG tracking, which the authors attributed to less expected pitch sequences (Keitel et al., 2023).

The differences in prediction accuracy between the baseline acoustical model (A) and the model with melodic expectation features (AM) allowed us to determine the melodic expectation enhancement effect (Δr). This effect was greater for the original music than for chimeric music. It was also greater for the full model (AM-A) than for either the pitch-only or time-only models (AMp-A and AMt-A, respectively).These differences demonstrate that chimeric music’s structural violations resulting from decoupling pitch and time indeed influenced melodic expectation encoding (**Figure 2d**).

The neural tracking (r) of both baseline model (A) and melodic expectation model (AM) was significantly correlated with the length of musical training, aligned with previous studies where musicians have stronger encoding of music (Di Liberto, Pelofi, Bianco, et al., 2020; Di Liberto, Pelofi, Shamma, et al., 2020). However, unlike Di Liberto, Pelofi, Bianco, et al. (2020), no correlation was found between musical training and the expectation enhancement effect (Δr).

Interestingly, there was no significant difference in TRF waveforms between original and chimeric music within the AM model, suggesting that while the effectiveness of expectation mechanisms may vary with the musical structures, the underlying neural encoding processes may remain consistent. A second explanation could be that the differences were small and distributed across the scalp, which would allow them to be seen in EEG prediction but not in the response waveforms at single electrodes.

In our analysis on the downbeat encoding with TRF, the note onset regressor had a TRF similar to the canonical morphology of the auditory evoked potential (Lalor et al., 2009; Picton et al., 1974), characterized by P1 and P2 components. This onset-evoked potential was the same for original and chimeric music. Further exploration of downbeat encoding with TRF revealed that downbeat notes elicit neural responses distinct from non-downbeats. This distinction was evident in both original music with the DBo regressor (which reflects the coupled downbeat pitch and time) and chimeric music with DBct regressor (which reflects downbeats defined by time). The neural downbeat response was enhanced around 185 ms, reflecting a larger P2 peak for downbeats.

These two results together suggest the distinct neural responses elicited by downbeat notes in comparison to non-downbeat notes, highlighting the significance of downbeats in the cognitive processing of musical structures. This is also in line with the previous study where the accented stimuli showed an enhanced N1/P2 complex in perception than unaccented within a percussive rhythmic pattern (Abecasis et al., 2009; Schaefer et al., 2011). It should be noted, however, that all notes in our stimuli, including downbeats, were of equal level. In other words, downbeats were defined entirely by their musical context.

One novel aspect of this study is the delineation of pitch structure’s impact on downbeat encoding. While the pitch component alone (Dcp) did not significantly affect the neural response to downbeats, the interaction between downbeat-pitch and downbeat-time term (Dcpt) showed a change in neural activity. This suggests that the cognitive processing of downbeats in music is multi-dimensional, where the interaction of the pitch and time structure of music both play a role. The impact of musicianship on neural encoding of music was demonstrated by a significant correlation between years of formal musical training and the P2 amplitude for note onsets. The P2 component is often associated with auditory attention (Picton & Hillyard, 1974) and sensitive to neuroplasticity (Atienza et al., 2002; Shahin et al., 2003; Tremblay et al., 2001), indicating that more musically trained individuals may exhibit enhanced processing of novel musical stimuli (Di Liberto, Pelofi, Shamma, et al., 2020; Shahin et al., 2003). However, unlike other studies that have reported differing rhythmic processing in musicians (Abecasis et al., 2009; Zhao et al., 2017), our results suggest that while musicianship appears to influence the neural encoding of basic auditory features such as note onsets, a uniform effect on the encoding of more complex musical structures such as downbeats may be harder to measure.

Another aspect of our results was the high correlation between the ABR waveforms generated by original and chimeric music, which aligns with the canonical ABR morphology, suggesting the subcortical level processing of auditory information was likely to be the same, regardless of the high level complexity of the musical stimuli (Shan et al., 2024).

Using chimeric music to decouple the two fundamental dimensions of melody—pitch and time— we confirmed that the structure of each is processed by listeners, and also found that their interaction is reflected in the listeners’ neural responses. This interaction plays a crucial role in developing the musical structure and pattern (Lerdahl, 2019; Tillmann, 2012), and thus impacts how music is perceived and processed by the human brain.

## Acknowledgments

Research reported in this publication was supported by NSF CAREER grant 2142612 and a grant from the Schmitt Program in Neuroscience.

