## supplemental materials for "Chimeric music reveals an interaction of pitch and time in electrophysiological signatures of music encoding"

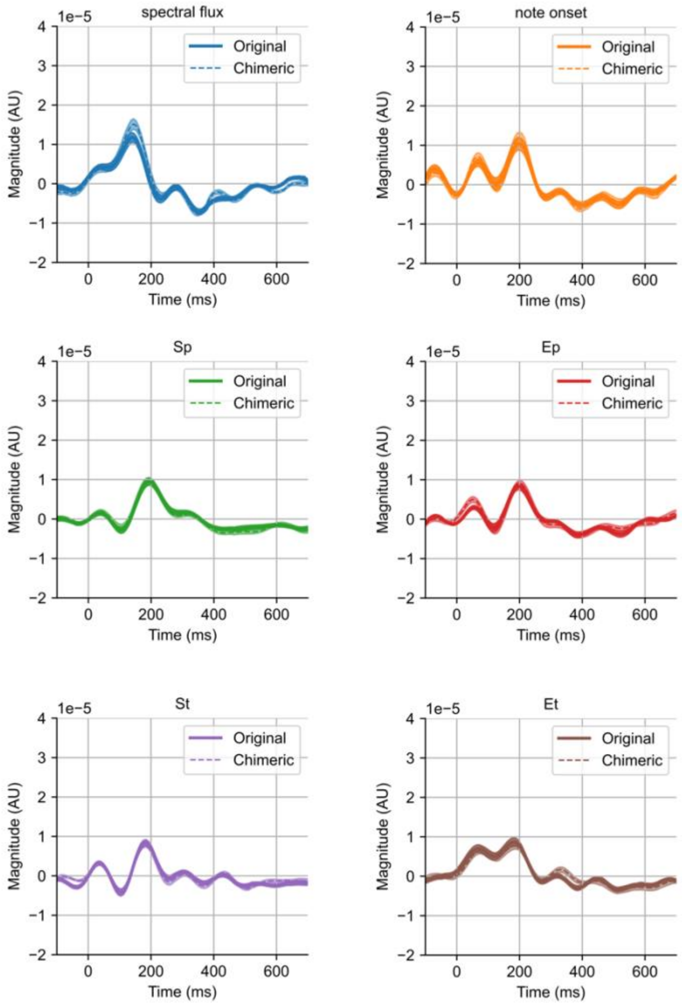


**Figure S1.** TRF weights from AM model at Fz channel. Each waveform is the weights from each regressor: spectral flux, note onset, Sp, Ep, St, Et from LTM and STM features. Shaded area shows ± 1 SEM. No significant difference was found between the waveform of the TRF derived from original and chimeric music.

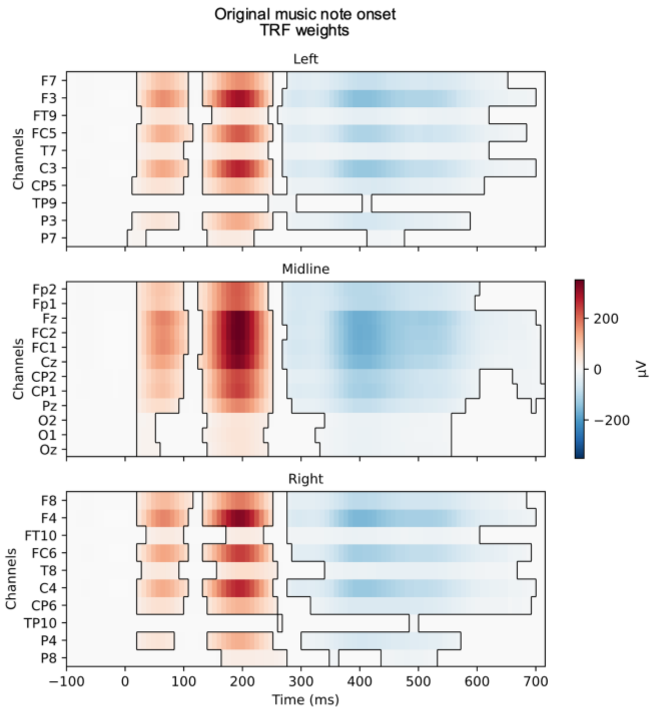


**Figure S2.** The TRF weights of original music note onset. Significant clusters of time points and channels is shown from permutation test.

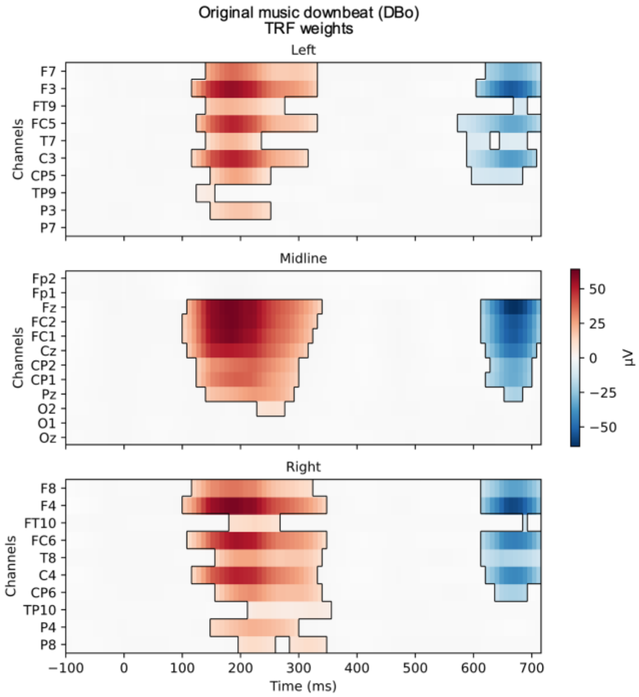

**Figure S3.** The TRF weights of original music DBo. Significant clusters of time points and channels is shown from permutation test.

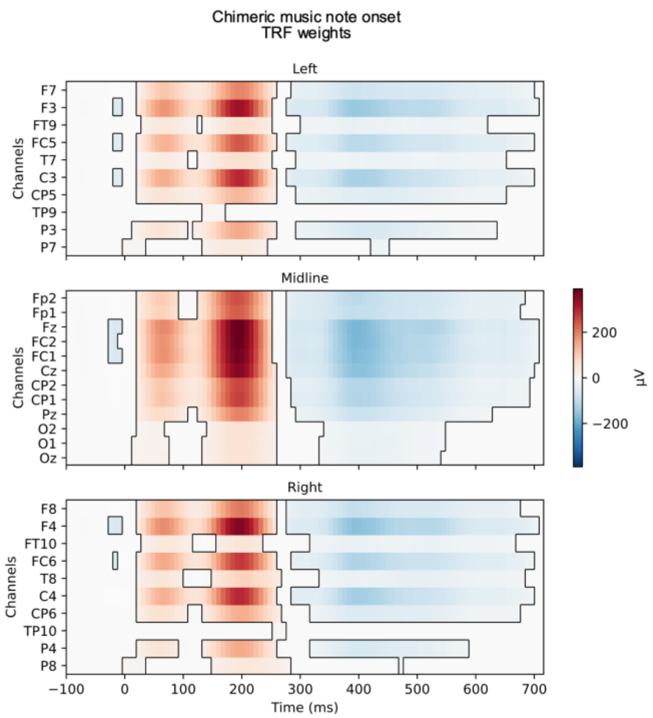


**Figure S3.** The TRF weights of chimeric music note onset. Significant clusters of time points and channels is shown from permutation test.

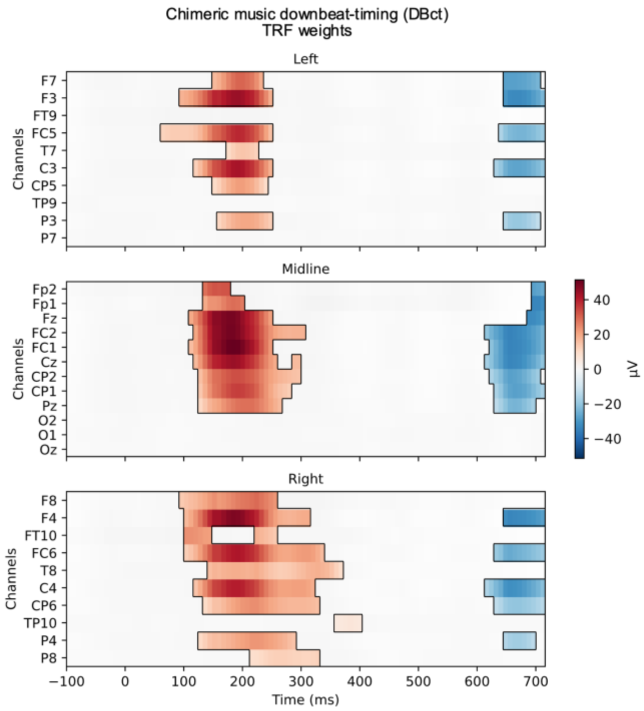


**Figure S4.** The TRF weights of chimeric music DBct. Significant clusters of time points and channels is shown from permutation test.

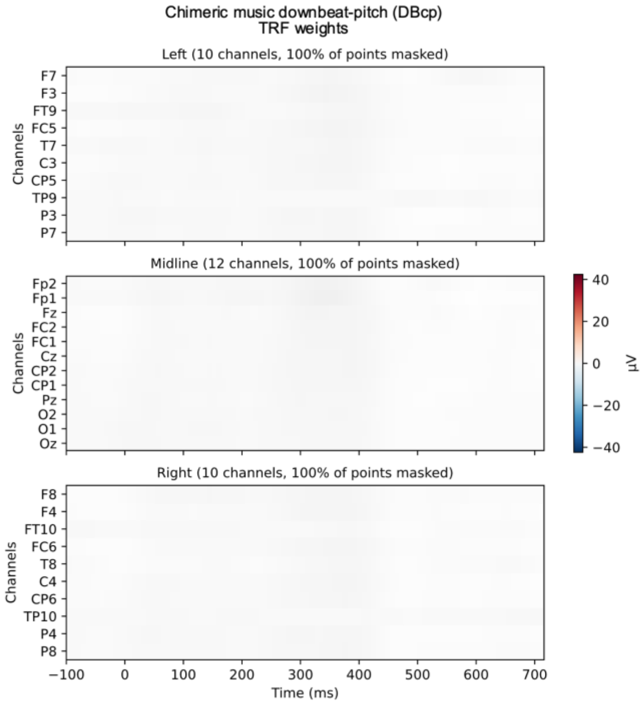


**Figure S5.** The TRF weights of chimeric music DBcp. There is no significant cluster found from permutation test.

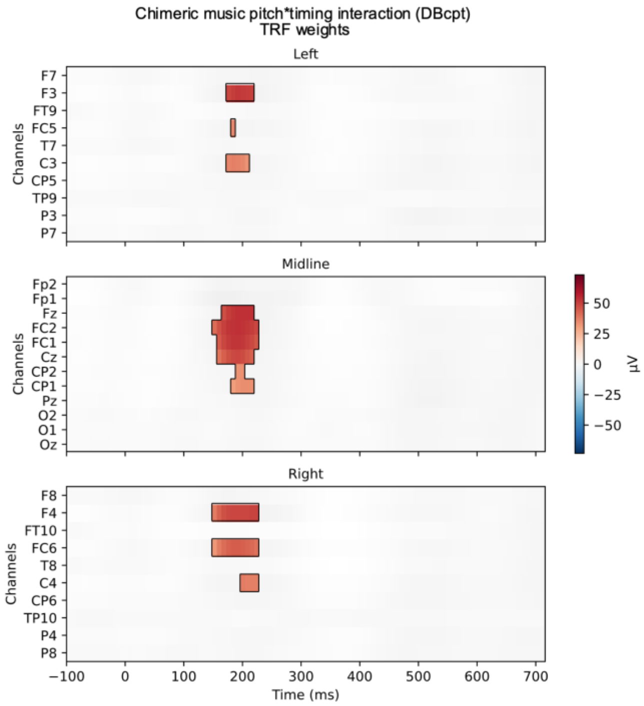


**Figure S6.** The TRF weights of chimeric music DBcpt. Significant clusters of time points and channels is shown from permutation test.
